# Physical models for chromosome organization to predict multi-contact statistics

**DOI:** 10.1101/2022.05.17.492279

**Authors:** Janni Harju, Joris J.B. Messelink, Chase P. Broedersz

## Abstract

Chromosome organization in both eukaryotes and prokaryotes is highly regulated. Organizing mechanisms, such as loop-extrusion, have been extensively studied using Hi-C methods, which measure pairwise contacts between chromosomal regions. New multi-contact methods additionally measure which chromosomal contacts occur simultaneously. Here, we develop three predictors of baseline multi-contact frequencies given pairwise contact data, corresponding to distinct physical limits, and argue that a comparison between data and prediction can lead to biological insight. We test these predictors for two simulated polymer models with cross-linking or loop-extrusion, and find that simulated three-point contacts are only predicted by the physically appropriate approximation. Finally, we apply our approach to previously published experimental multi-contact data from human chromosomes. Strikingly, we discover that observed three-point contact frequencies are well predicted by a formula based on loop-extrusion, suggesting that multi-contact data can give insight into chromosome organization mechanisms.

---

Both eukaryotic and prokaryotic chromosomes are compressed by orders of magnitude, while remaining accessible to replication and transcription machinery. This functional organization of chromosomes can be studied using chromosomal capture methods, such as Hi-C, which track the frequency at which pairs of chromosomal loci are spatially proximate, or “in contact” [1–3]. However, since traditional Hi-C methods provide only population-averaged pairwise contact counts, new methods must be used to assess whether chromosomal contacts occur simultaneously. Patterns of simultaneous contacts could reveal the presence of epicenters of chromosomal activity, such as transcription factories [4] or super-enhancers [5]. In addition, simultaneous contacts could offer insight into interactions between loop-extruding proteins that play a prominent role in chromosome organization, such as SMC complexes condensin and cohesin [6, 7].

To measure the frequencies of simultaneous contacts, several experimental protocols have been implemented, yielding so-called “multi-contact data”. The most general of these methods are single-cell contact-tracking experiments, such as super-resolution imaging [8–10], single-cell Hi-C [11, 12], or single-cell SPRITE [13]. Others have adapted chromosomal capture methods to study the average frequency of simultaneous contacts between three or more sites [14–19]. Although these new experimental methods hold promise for teaching us more about chromosome organization, it remains challenging to quantitatively analyze multi-contact data.

What can we learn from the frequency of a set of simultaneous chromosomal contacts? One approach is to compare the multi-contact frequency to an expectation value based on pairwise contact frequencies. Such a comparison tells us whether chromosomal contacts appear to be mutually enhanced or correlated, which can provide clues about the underlying chromosome organization. However, it is unclear what type of prediction should be used as a baseline, since even contacts on a non-interacting polymer are non-trivially correlated [20]. Due to this ambiguity, experimental multi-contact data have been compared to a wide range of predictions, leading to varying conclusions about to what extent chromosomal contacts are correlated [8, 15, 19, 21].

Here, we propose that different multi-contact predictions based on pairwise frequencies can be motivated by distinct physical models for baseline chromosomal contact formation. We discuss the appropriate predictions for an ideal polymer, a polymer with active loop-extruders, and a polymer with pairwise interactions between monomers. We test these predictions against multi-contact data from simulations with either random cross-linking between nearby monomers, or with loop-extruding condensins. The comparison shows that only the appropriately motivated approximation is predictive of multi-contact data. Finally, we compare the performance of all three approximations for previously published data for human IMR90 cells [8]. We find that the multi-contact data is accurately predicted by the loop-extruder approximation, thereby illustrating how our approach can be used to interpret multi-contact data.

## Theory

Currently, three-point contact frequencies are the most prevalent type of multi-contact data available [14–19] (Fig. 1(a)). To investigate how these data can be analyzed, we start by introducing predictions for three-point contact frequencies in three distinct physical scenarios, and briefly discuss how these approaches can be adapted for other types of multi-contact data.

**FIG. 1.**
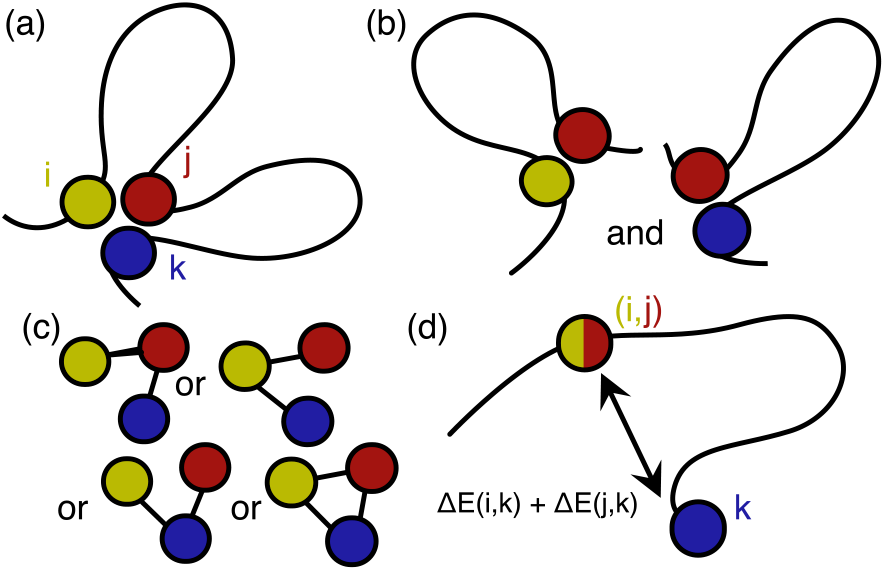
Visualization of approximations for three-point contact frequencies. **(a)** We consider a three-point contact between sites *i* < *j* < *k*. **(b)** Ideal polymer approximation. Here, the probability of the three-point contact is the probability of the two smallest loops (*i,j*) and (*j, k*). **(c)** Loop-extruder approximation. Now the probability of a three-point contact is given by the probability of at least two loop-extruders being present. **(d)** Pairwise interaction approximation. In this case, the three-point contact (*i, j, k*) is equivalent to a contact (*k*, (*i,j*)), with an effective length |*j – k*| and interactions between the neighbourhoods of k and the other two points.

As a simple starting point, we consider contacts on a non-interacting ideal polymer in a cell, equivalent to a confined random walk. The contact probability *P_0_*(*i,j*)between any two sites *i, j* on the polymer is directly determined by the genomic length |*i – j*| between them.

When two contacts form simultaneously, say between *i, j* and *j,k* with *i* < *j* < *k* (Fig. 1(b)), the contacts corresponding to the two smallest loops (i,j) and (j,k) are statistically independent. Hence, in this non-interacting limit, the three-point contact probability is given by

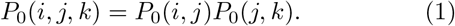

By replacing the non-interacting pairwise contact probabilities *P_0_*(*i,j*) on the right hand side with those found experimentally for a chromosome, *P*(*i,j*), this formula provides a simple approximation for three-point contact probability *P*(*i, j, k*) that aims to take into account some of the chromosome structure present *in vivo*, e.g. variations in effective stiffness along the chromosome [22]. This prediction can also be adapted to describe other types of multi-contact statistics by decomposing a multicontact structure onto a set of loops that are independent on an ideal polymer [20]. We hence refer to this approximation as the *ideal polymer approximation*.

A key organizational mechanism in both pro- and eukaryotic chromosomes is active loop-extrusion [23, 24]. Therefore, we next consider the limit where noninteracting loop-extruders *dominate* chromosomal contact formation. Loop-extruder motor proteins, such as condensins, can load onto a strand of DNA and then actively traverse in one dimension along DNA to extrude a loop [25, 26]. Thus, we assume that their positions are independent of the three-dimensional configuration of the polymer. Importantly, this implies that, unlike ideal polymer contacts, loop-extruder-mediated contacts occur largely independently of each other, since the presence of a contact elsewhere on the polymer is expected to have little impact on the procession of a loop-extruder.

Let *P*_LE_(*i, j*) be the probability that the loci i and j are linked by a loop-extruder. The probability that there are at least two loop-extruders between sites *i, j, k*, resulting in a three-point contact, is then given by (Fig. 1(c))

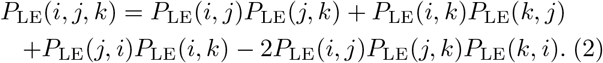

The last term ensures that the case with three loop-extruders is not over-counted. When chromosomal contacts are dominated by loop-extruders, this formula can be used to approximate three-point contact probabilities *P*(*i, j, k*), by substituting all P_LE_’s on the right hand side by the respective pairwise contact probabilities. We note that Eq. (2) is related to an existing algorithm not motivated by loop-extrusion [15] (see Supplemental Material at [27] for a discussion). Our approach can be extended to other types of multi-contact statistics by considering all possible combinations of loop-extruders that would explain the set of contacts. We hence refer to Eq. (2), and its extensions for other multi-contacts, as the *loop-extruder approximation*. Since this approximation ignores how a contact brings together two polymeric regions, and hence enhances contacts between them, it is unclear how accurately Eq. (2) could predict chromosomal multi-contact data.

Given the simplifications made by the ideal polymer and loop-extruder approximations, we next consider a third scenario that takes into account both polymeric effects and affinities of chromosomal regions for each other, e.g. as induced by bridging proteins or other agents effectively linking two sections of the chromosomes [28]. Specifically, we model chromosomes as equilibrium polymers with close-range interactions between chromosomal loci i and j [29]. Such a model can also be motivated by information theory: given pairwise contact frequencies, the least assuming—or maximum entropy—distribution of chromosome configurations is mathematically equivalent to a polymer with close-range pairwise interactions [30, 31].

Let **X** = {**x_i_**}_iϵ{1,2,…N}_, be the configuration of a polymer with *N* monomers. For simplicity, we consider a coarse-grained lattice polymer representation, where the lattice spacing is of the order of the interaction length, such that a configuration has a probability

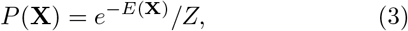

where 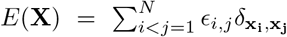 is the energy of the configuration in units of *k_B_T*, with δ_xi_,_xj_ the Kronecker delta-function, and *Z* is the partition function. The probability of a contact (*i, j*) can be expressed as

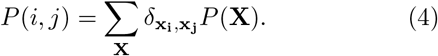

To derive an approximation for multi-contact statistics, note that the probability of a three-point contact (*i, j, k*) is determined by its energetic and entropic cost

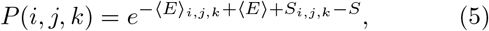

where ⟨*E*⟩ is the expected value of the configuration energy, *S* = – ∑_X_ *P*(**X**) log(*P*(**X**)) is the configuration entropy, and is a conditional average over configurations with this three-point contact.

When contact energies are additive (Δ*E_i,j,k_* ≈ Δ*E_i,j_* + Δ*E_j,k_* + Δ*E_k,i_*) and sufficiently weak (Δ*S* ≈ Δ*S_0_*), we can approximate the probability of a three-point contact as (see Supplemental Material at [27] for derivation)

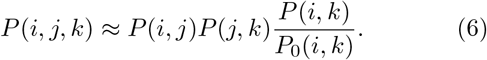

The terms *P*(*i,j*) and *P*(*j, k*) include the energetic and entropic costs of the two loops in the three-point contact, whereas the term 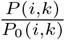 adds an approximation for the energetic cost of contact (*i, k*) (Fig. 1(d)). The formula can be extended for other types of multi-contact statistics by multiplying the ideal polymer approximation with a factor of *P/P_0_* for every contact not already considered. However, the error is expected to grow as more energies are estimated via *P/P_0_*. Nevertheless, this pairwise interaction approximation takes into account both the energetic cost of the largest loop, unlike the ideal polymer formula, and its reduced entropic cost due to the other two loops, unlike the loop-extruder formula.

## Testing approximations against simulated data

To test whether our three approximations are capable of predicting three-point contact frequencies in their associated physical limits, we examine simulated three-point contact data from a model of loop-extruding condensins on a bacterial chromosome [7] (Fig. 2(b)), and a model with cross-linkers randomly binding spatially proximal regions of a polymer (Fig. 2(a)). In the condensin simulations, we consider a set of 40 loop-extruders that load onto a circular chromosome primarily at a single site (see Supplemental Material at [27] for details), giving rise to a Hi-C map with an off-diagonal line of contacts (Fig. 2(c)), similar to bacteria such as *Caulobacter crescentus* [3] or *Bacillus subtilis* [32]. We expect alternative simulation schemes for loop-extrusion [33, 34] to yield similar results, as long as the loop-extruders mainly move in one dimension along the chromosome, and only interact weakly. In the the cross-linker simulations, we define a binding affinity for each polymer site, *ϵ_i_*, such that the energy of a contact (*i,j*) is *ϵ_i_* + *ϵ_j_*. As simple test cases, we consider affinities that vary as a cosine function over the genomic position, resulting in a check-patterned Hi-C map (Fig. 2(c)), as well as affinities that are randomly sampled independent of genomic position (Supp. Fig. S1). The loop-extruder and cross-linker simulation models differ qualitatively in the way in which contact formation is induced, allowing us to test whether different multicontact predictors work better in different limits.

**FIG. 2.**
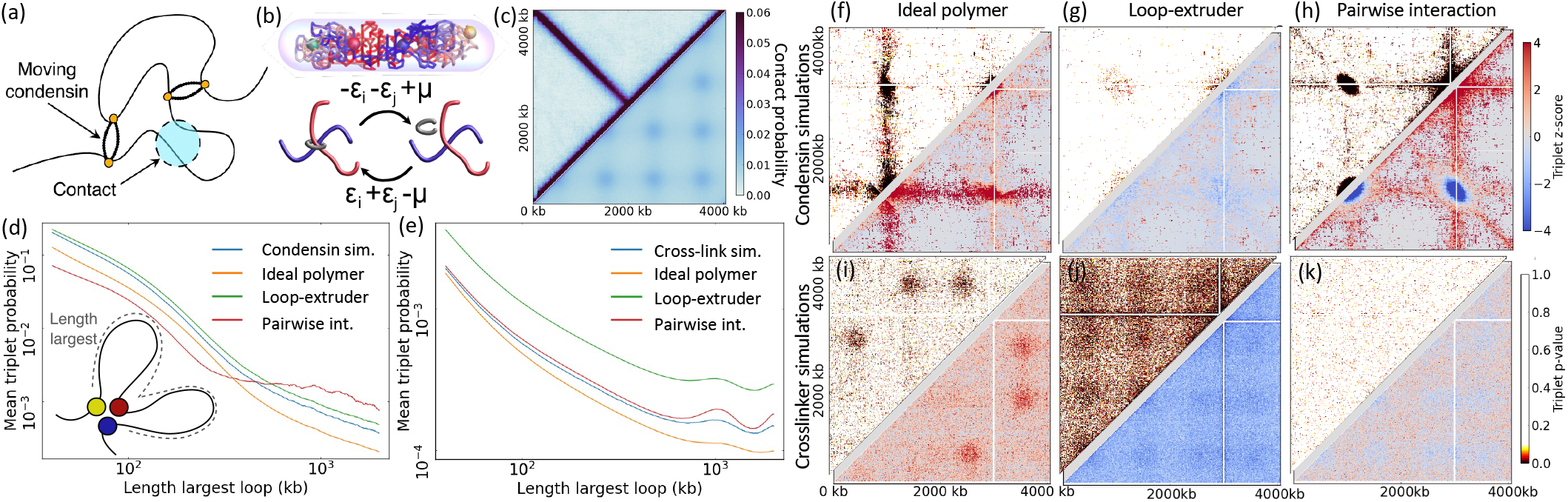
Cross-linker and condensin simulations. **(a)** Condensin simulations [7, 35] are conducted by updating the locations of moving condensins on a 3D bead-spring polymer, and letting the polymer relax given the constraints. Contacts are defined as loci within a predefined range of 5 simulation units. **(b)** Cross-linker simulations are conducted in a lattice polymer set-up, using the dimensions of a *C. crescentus* chromosome [31]. For a polymer with *N* monomers, we define a chemical potential *μ* = 5*k_B_T* and a binding affinity *ϵ_i_* = cos(8*πi/N*)*k_B_T* for cross-linkers, giving ⟨n_CL_⟩ = 41.3. **(c)** The simulated contact probability maps for condensin (top-left) and cross-linker (bottom-right) simulations. **(d-e)** Plots show *P_3_(s*) curves for data sampled from the condensin and cross-linker simulations, and predictions calculated using pairwise contact frequencies (see Supplemental Material at [27] for simulation details). **(f-k)** Comparison of the condensin and cross-linker simulation data to the approximations. Top-left: *p*-values calculated by presuming the three-point contact data is binomially distributed with a probability calculated based on an approximation. Bottom-right: Z-values calculated for the three-point contact map compared to predictions. All plots are constructed for the bait point corresponding to 3000 kb.

To visualize how our three-point contact frequency predictions perform, we define *P_3_(s*) as the average probability of three-point contacts where the largest loop has genomic length s (see Supplemental Material at [27] for details). We find that s is more indicative of prediction performance than the size of either smaller loop in a three-point contact (Supp. Fig. S2). In addition, to avoid averaging over genomic location, we follow [14, 15, 18] and visualize three-point contact probabilities for a given bait point k as heatmaps where the intensity at point (*i, j*) is the frequency of three-point contacts (*i,j, k*).

The three-point contact frequencies of the condensin simulations are best predicted by the loop-extruder approximation (Fig. 2(g)), for a range of bait points (Supp. Fig. S3). This approximation gives only a slight overestimate for loops of all sizes (Fig. 2(d)), and after correcting for multiple hypothesis testing, few *p*-values are significant (Supplemental Figure S4). The ideal polymer formula, by contrast, fails to predict several lines on the three-point contact map (Fig. 2(f)). These missing lines correspond to contact triplets where the largest loop—ignored by the ideal polymer approximation—lies on the condensin trajectory. The pairwise interaction formula is also inaccurate, partially because it presumes simultaneous interactions between all three points. We conclude that the simulated three-point contact data is accurately predicted by the loop-extruder approximation, reflecting that the model’s contact formation is predominantly driven by the localization of weakly interacting condensins.

The *P_3_(s*) curve (Fig. 2(e)) of the cross-linker model, by contrast, is best predicted by the pairwise interaction formula. The three-point contact map (Fig. 2(k)) shows that the approximation slightly under-estimates the frequencies of the most-likely three-point contacts. When multiple hypothesis testing is accounted for, most deviations from the pairwise interaction formula do not appear significant (Supp. Fig. S4). By contrast, the ideal polymer and loop-extruder formulas give significant under- and overestimates, respectively. Results are similar for different values of the cross-linker chemical potential μ, and when random binding affinities are considered, but with deviations appearing for large numbers of bound cross-linkers (Supp. Fig. S1). We hence conclude that when cross-linking is sufficiently weak, the pairwise interaction approximation gives an accurate prediction for our cross-linker simulation data.

So far, we have used absolute contact frequencies to predict three-point contact statistics. However, most chromosomal capture methods only yield relative contact counts, which scale with actual contact probabilities. If relative counts are used, multi-contact frequencies can only be predicted up to a constant prefactor, which can be set by scaling predictions to match the measured total number of three-point contacts [15, 19]. Suprisingly, we find that even if an approximation is inaccurate when predicting absolute three-point contact frequencies, it can perform well after such a scaling (Supp. Fig. S5). This indicates that we should be cautious when using relative (multi-)contact frequencies: the best predictor of our simulated multi-contact data only reflected the dominant contact formation mechanism for the system when absolute contact frequencies were used.

## Experimental multi-contact data

Having shown that simulated multi-contact data can be predicted by simple approximations that use pairwise contact frequencies, we next apply our approach to experimental multi-contact data. Specifically, we analyze the super-resolution chromatin tracing data for human IMR90 cells from Ref. [8] (see Supplemental Information [27] for details). These data can be used to find the absolute frequencies of pairwise contacts and three-point contacts (Fig. 3(a)), avoiding the need for scaling predictions, which we found to distort predictions for simulated data. The authors previously tested for ‘‘cooperative, higher-order chromatin interactions” [8] using the hypothesis *P*((*j, k*)|(*i, j*)) = *P*(*j,k*), related to the ideal polymer approximation Eq. (1), and found significant three-point contact enhancement compared to this formula. We now ask whether the loop-extruder or pairwise interaction approximations can better predict the multi-contact data.

**FIG. 3.**
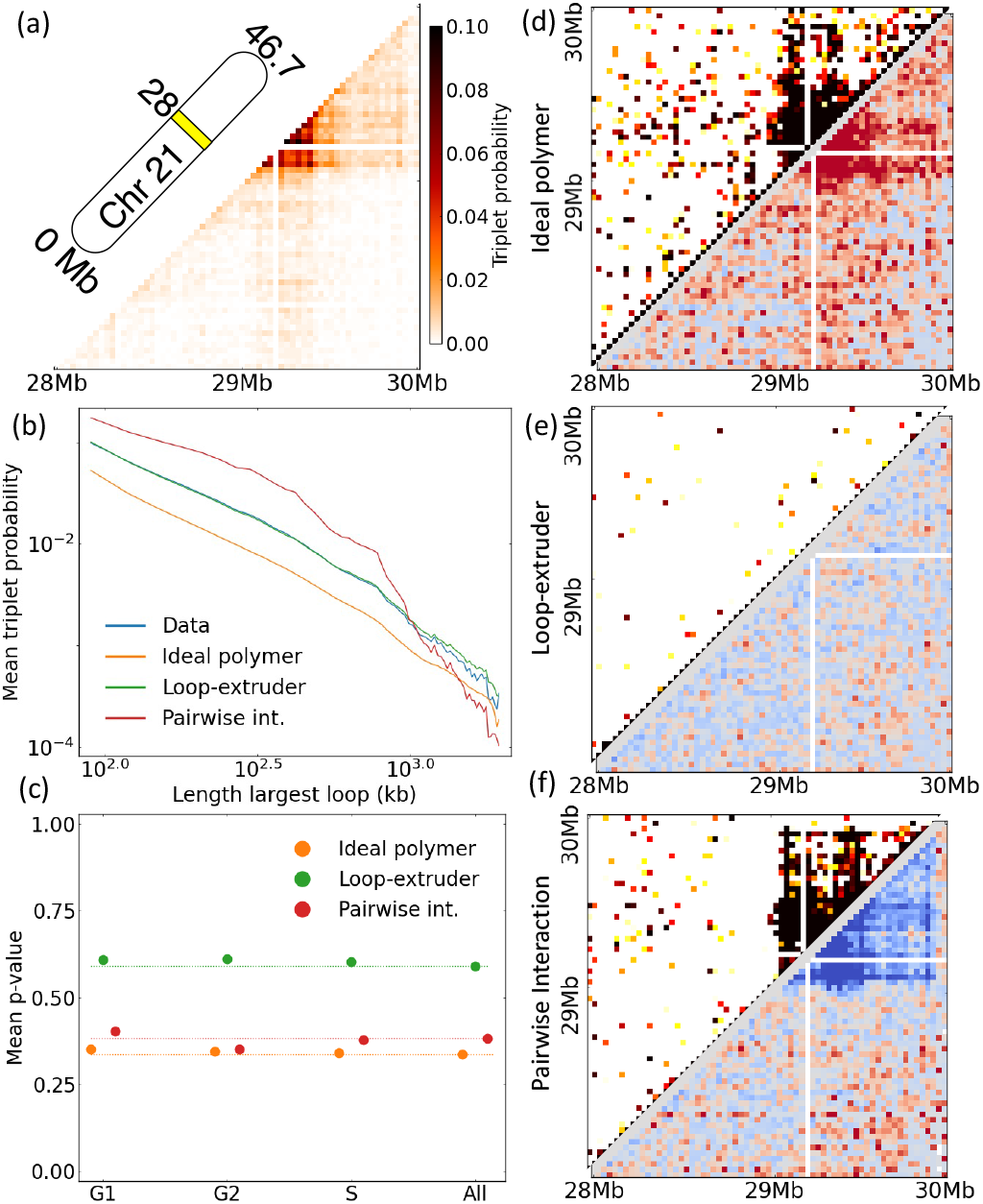
Three-point contact predictions applied to experimental data on human chromosomes from Bintu *et al*. **(a)** The three-point contact probabilities for an arbitrary bait point located at 29.1 Mb on chromosome 21. **(b)** Similar to (Fig. 2(d,e)). Blue line describes the experimentally found contact triplet probabilities for chromosome 21 of 1574 IMR90 cells in the G1, G2 or S phase. The predicted curves were calculated using the experimentally found pairwise contact probabilities. Contacts were defined as spatial separations of < 150 nm between the centers of labelled 30 kb regions. **(c)** Comparison of the *p*-value averaged over three-point contacts for data divided by cell cycle phase or combined together. Dashed lines indicate the *p*-values for the combined data. 288 samples were used to calculate all *p*-values, to ensure that results are comparable. **(d-f)** p-value and *z*-score plots for the ideal polymer, loop-extruder, and pairwise interaction formula, color-bars in (Fig. 2(h,k)).

For simplicity, we first focus on data averaged over the S, G1, and G2 cell cycle phases. Strikingly, the loop-extruder approximation performs by far the best in describing the experimental multi-contact data (Fig. 3(b,e)), and at an accuracy comparable to our results for condensin simulations (Fig. 2(d,g)). Indeed, when controlling for multiple hypothesis testing, hardly any deviations from the loop-extruder formula can be considered significant (Supp. Fig. S6). Consistent with the earlier analysis [8], the ideal polymer approximation gives an underestimate of three-point contact frequencies (Fig. 3(b,d)). The pairwise interaction formula, on the other hand, gives persistent overestimates for three-point contacts within TADs (Fig. 3(f), Supp. Fig. S7). These overestimates could arise for the same reason as observed in our condensin simulations (Fig. 2(f)); if a three-point contact occurs due to a collision between two loop-extruders, there is no “attractive force” associated with the third contact, as presumed by the pairwise interaction formula. We thus conclude that the three-point contact data is largely predictable from pairwise contact frequencies, using an approximation where contacts are dominantly caused by weakly interacting loop-extruders.

To test whether the change of the pairwise contact probabilities throughout S, G1 and G2 phase has an impact on the quality of our approximations, we repeated the analysis for contact data separated by cell cycle stage. This separation of data does not noticeably improve the mean p-value for our predictions (Fig. 3(c)). We hence conclude that contact correlations due to loops being enhanced during the same cell cycle stage are not significant at this sample size.

As an alternative approach, we repeated our analysis for these experiments using the three-point contact frequency prediction recently proposed in Ref. [21]. Their method is based on inferring a set of spring constants from Hi-C data, and is hence computationally more complex than the loop-extruder or ideal polymer approximations, which calculate a set of three-point contact frequencies with optimal N^3^ scaling. Nevertheless, for the Bintu *et al*. data analyzed, we found that the loop-extruder formula performed slightly better than the spring model [21] (mean *p*-values 0.59 vs. 0.52; (Supp. Fig. S8)). Furthermore, unlike the loop-extruder prediction, the spring model prediction had to be scaled to match the observed number of three-point contacts, which can can lead to seemingly better prediction performance (Supp. Fig. S5). The comparatively good performance of our loop-extruder approximation suggests that our simple approaches can provide useful benchmarks when constructing more complicated predictions for multi-contact frequencies.

## Discussion

In summary, we physically motivated three predictions for multi-contact data: the loop-extruder, the ideal polymer, and the low-energy pairwise interaction approximations. When we tested these approximations against data simulated with a model with weakly interacting loop-extruders and a model with chromosome cross-linkers, we found that the approximation scheme associated with the dominant mechanism of contact formation performed well in predicting three-point contact statistics. When we applied our approach to published experimental data for IMR90 chromosomes [8], we found that only the loop-extruder approximation was capable of accurately predicting the data at the megabase scale. This suggests that loop-extrusion can have a measurable impact on chromosomal contact statistics, and that multi-contact data could hence be used to study mechanisms of chromosome organization.

We thank Hugo Brandāo for discussions and help with condensin simulations, and Pedro Olivares-Chauvet for help accessing experimental data.

