## Supplemental Information for "Physical models for chromosome organization to predict multi-contact statistics"

#### S1 Mathematical formulation of sampling method used by Olivares-Cadete *et al.*

We briefly explain how the sampling scheme introduced in [1] can be expressed as a mathematical formula. The authors consider three-point contacts $(i, j, k)$ for a given bait point $k$. They compare their experimental data to a sampled data set generated as follows:

1. Sample $i$ using the 4C data around the bait $k$ as weights. This corresponds to the probability $P(i|k) = \frac{P(k,i)}{\sum_m P(m,k)}$, the probability of a contact between $k$ and $i$, given that $k$ has a contact.
2. With a probability $p$, sample $j$ from the Hi-C neighbors of $i$ (probability $P(j|i)$). Else, sample $j$ from the 4C neighbors of $k$ (probability $P(j|k)$).

The authors pick the parameter $p$ by analyzing at what ratio the third point in their data set is closer to $i$ than to $k$. However, the same value that they arrive at, $p = 2/3$, can be mathematically justified. The sampling weight of the above algorithm is:

$$P(i, j, k) \propto P(i|k)[pP(j|i) + (1 - p)P(j|k)] + P(j|k)\left[pP(i|j) + (1 - p)P(i|k)\right] \tag{S1}$$

which can be reorganized as

$$P(i, j, k) \propto 2(1 - p)P(i|k)P(j|k) + pP(j|i)P(i|k) + pP(i|j)P(j|k). \tag{S2}$$

Requiring that all three terms have equal weights yields $p = 2/3$, and we then find

$$P(i, j, k) \propto P(j|i)P(i|k) + P(i|j)P(j|k) + P(i|k)P(j|k). \tag{S3}$$

This approximation differs from (Eq.2) in two ways: firstly, (Eq. S3) uses relative contact probabilities and gives a prediction up to a constant; secondly, the higher-order term of (Eq.2) is absent. The use of relative contact probabilities means that the prediction can be calculated using relative contact counts from Hi-C experiments. We note that when relative contact counts are used, however, the higher-order term from (Eq.2) cannot be included, since it scales as $P^3$ rather than $P^2$.

#### S2 Derivation of the pairwise interaction formula

Here we provide a derivation of the low-energy pairwise interaction formula, aimed at predicting three-point contact probabilities when we can associate an energetic cost with contact formation. For simplicity we consider a lattice-polymer representation, where contacts correspond to monomers occupying the same lattice site, and a contact between sites $(i, j)$ has an associated energy $\epsilon_{i,j}$. If a freely-jointed chain or bead-spring polymer representation was used, the delta function in Equation 5 would be replaced by a Heaviside step-function for the distance between two monomers. The following analysis is independent of this choice of representation, provided that a three-point contact is defined as pairwise contacts between all three sites.

To start, we define the model's effective free energy as $F = -\log(Z)$, where $Z$ is the partition function, defined as a sum over the statistical weights of all possible polymer configurations:

$$Z = \sum_{\mathbf{X}} e^{-E(\mathbf{X})}. \tag{S4}$$

The equality $F = \langle E \rangle - S$ holds, where $\langle E \rangle = \sum_{\mathbf{X}} E(\mathbf{X})P(\mathbf{X})$ is the expectation value of the energy, and $S = -\sum_{\mathbf{X}} \log(P(\mathbf{X}))P(\mathbf{X})$ is the distribution entropy.

To express probabilities of simultaneous contacts, we also define a class of constrained partition functions

$$Z(i,j) = \sum_{\mathbf{X}} e^{-E(\mathbf{X})}\delta^3(\mathbf{x_i} - \mathbf{x_j}), \tag{S5}$$

and their corresponding free energies $F(i,j) = -\log(Z(i,j))$, which can easily be generalized to higher-order multi-contacts. These tools allow us to write contact probabilities as

$$P(i,j) = e^{-F(i,j)+F} = e^{-\langle E\rangle_{i,j} + \langle E\rangle + S(i,j) - S}, \tag{S6}$$

and similarly for three-point contacts. Contact (set) probabilities can hence be expressed in terms of their expected energetic cost, here $\Delta E(i,j) := \langle E\rangle_{i,j} - \langle E\rangle$, and their entropic cost, here $\Delta S(i,j) := S(i,j) - S$. Finding an estimate for the probability of a set of contacts $\Gamma$ can hence be thought of as finding estimates for $\Delta E(\Gamma)$ and $\Delta S(\Gamma)$.

We first consider the energetic cost of a three-point contact:

$$\Delta E(i,j,k) = \sum_{(n,m)} \epsilon_{n,m}(P((n,m)|(i,j,k)) - P(n,m)). \tag{S7}$$

In the case where all contacts $(n,m)$ are strongly correlated with only one of the contacts $(i,j)$, $(j,k)$ or $(i,k)$, the right-hand side is approximated by $\Delta E(i,j) + \Delta E(j,k) + \Delta E(i,k)$. We note that this does not hold when, for example, loci $i$ and $j$ are close to each other.

To obtain an estimate for the entropic cost, we focus on the limit of small energies $\epsilon_{n,m}$, or equivalently high temperatures. In this limit, the system starts to resemble a non-interacting polymer, with contact probabilities $P_0(n,m)$. For the non-interacting system the ideal polymer formula for three-point contacts is exact, and hence

$$\Delta S_0(i,j,k) = \Delta S_0(i,j) + \Delta S_0(j,k). \tag{S8}$$

When interaction energies are sufficiently small, a similar factorization will hold, since the entropic cost of a combination of contacts is well approximated by its entropic cost on the non-interacting polymer.

To evaluate how $\Delta S$ approaches $\Delta S_0$, we use the cumulant expansions for the free energy differences [2]:

$$F - F_0 = -\sum_{n=1}^{\infty} \frac{(-1)^n \kappa_n}{n!}, \tag{S9}$$

$$F(\Gamma) - F_0(\Gamma) = -\sum_{n=1}^{\infty} \frac{(-1)^n \tilde{\kappa}_n}{n!}, \tag{S10}$$

where $\kappa_n$ is the $n$th cumulant of the total energy $E$ calculated for the non-interacting polymer, and $\tilde{\kappa}_n$ is the $n$th cumulant of $E$ calculated on the non-interacting polymer with the contact(s) in the set $\Gamma$ constrained.

Using S6, we hence have that

$$\log\left(\frac{P(\Gamma)}{P_0(\Gamma)}\right) = F - F_0 - F(\Gamma) + F_0(\Gamma) = \sum_{n=1}^{\infty} \frac{(-1)^n(\tilde{\kappa}_n - \kappa_n)}{n!}. \tag{S11}$$

We next define a similar expansion for $\Delta E(\Gamma)$. We first introduce an inverse temperature $\beta$, writing $F = F(\beta = 1)$, with $F(\beta) = -\log(\sum e^{-\beta E})$. In this case,

$$\Delta E(\Gamma) = \frac{\partial}{\partial \beta} F(\Gamma)|_{\beta=1} - \frac{\partial}{\partial \beta} F|_{\beta=1} = -\sum_{n=1}^{\infty} \frac{(-1)^n(\tilde{\kappa}_n - \kappa_n)}{(n-1)!}, \tag{S12}$$

where we included factors of $\beta^n$ in Equations S9 and S10 and differentiated. Note that the cumulants $\kappa_n$ and $\tilde{\kappa}_n$ are expectation values of polynomials of the energy on the non-interacting polymer, and hence independent of $\beta$.

Adding expression S11 and S12 gives

$$\log\left(\frac{P(\Gamma)}{P_0(\Gamma)}\right) + \Delta E(\Gamma) = -\sum_{n=2}^{\infty} \frac{(-1)^n(\tilde{\kappa}_n - \kappa_n)}{(n-1)!}\left(1 - \frac{1}{n}\right). \quad \text{(S13)}$$

Using that the leading order term in S12 is $\Delta E_0(\Gamma)$, we see that the right-hand-side of S13 is of order $\frac{\Delta E - \Delta E_0}{2}$, provided that the cumulants of order $n > 2$ are small. This limit is approached when the interaction energies are sufficiently small, or when contacts on the non-interacting polymer are sufficiently independent that a generalized central limit theorem applies for the sum $E = \sum_{(n,m)} \epsilon_{n,m}\delta^3(\mathbf{x_n} - \mathbf{x_m})$.

We have hence shown that

$$\Delta S(i,k) - \Delta S_0(i,k) = \log\left(\frac{P(i,k)}{P_0(i,k)}\right) + \Delta E(i,k) = \mathcal{O}\left(\frac{\Delta E(i,k) - \Delta E_0(i,k)}{2}\right). \quad \text{(S14)}$$

The correction term is of order $\epsilon(P - P_0)$. In the limit of small energies, when $\epsilon_{i,j}(P(i,j) - P_0(i,j)) << 1$, the entropic cost of contacts is hence approximated by their entropic cost on the non-interacting polymer. This implies that the entropic cost for multi-contacts approximately factorizes. Furthermore, we have that the energetic cost of a contact, $\Delta E(i,j)$, can be approximated by $-\log(P(i,j)) + \log(P_0(i,j))$.

Using these results, we find that in the case when $i, j, k$ are sufficiently separated so that their energetic costs add, and when interactions are sufficiently weak such that $1/2|\Delta E(i,k) - \Delta E_0(i,k)| << 1$, we expect that

$$P(i,j,k) \approx e^{-\Delta E(i,k)} P(i,j)P(j,k) \approx \frac{P(i,k)}{P_0(i,k)} P(i,j)P(j,k), \quad \text{(S15)}$$

as was to be demonstrated.

### S3 Condensin simulations

The script used for [3], available at [4], was adapted as follows; a single condensin loading site at 1 kb was used, and condensins were assumed to move at equal speeds in both directions on the chromosome (parameter "wind" set to zero in simulations). Raw configuration data and Julia scripts for analysis are available in [5]. Contacts were defined as a distance of less than 5 simulation units between monomers, and three-point contacts as events where at least two contacts were present between three monomers. We note that using this definition, the pairwise interaction approximation makes a further approximation; it neglects the fact that two loci in a three-point contact can be more than a cross-linking radius apart, which can change both the entropic and energetic cost of this secondary contact. Contact and three-point contact frequencies were calculated by sampling 3000 polymer configurations.

### S4 Cross-linker simulations

Cross-linker simulations were adapted from a coarse-grained lattice polymer model for the *C. crescentus* chromosome [6], which is available at [7]. In short, we used a Metropolis Monte Carlo simulation for a circular lattice polymer in a cylindrical volume of confinement, with the polymer length and volume size constrained by imaging data. For a given set of affinities $\epsilon_i$ and a chemical potential $\mu$, after each (attempted) monomer move, we picked a random monomer and looped through its cross-links, detaching each cross-link with a probability $P_{\mathrm{UB}} = 1/2$. We then picked another monomer $i$, looped through all monomers $j$ on the same lattice site, attempting cross-linking with a probability $P_{\mathrm{UB}}e^{-(\epsilon_i+\epsilon_j-\mu)}$. Note that for all simulations $\epsilon_i + \epsilon_j - \mu \geq 1\mathrm{k_B T}$, and hence all binding probabilities were smaller than 1. Cross-linked monomers were not allowed to move.

#### S5 Data from Bintu *et al.*

We analyzed super-resolution chromatin tracing data published by Bintu *et al.* [8]. The authors imaged 65 neighboring 30 kb intervals of human chromosome 21, and thus gathered data on their relative positions. The data files containing relative positions of loci were used to calculate the frequencies of contacts (two loci separated by a distance less than 150 nm) and three-point contacts (three loci with at least two contacts between them), using a Julia script available in [5].

#### S6 Non-interacting simulations

The pairwise interaction formula (Eq.6) requires the probabilities of contacts on a non-interacting, ideal polymer, with the same length and confinement volume as the chromosome.

For the cross-linking model, the simulations were run with no cross-links. For the loop-extrusion model, excluded volume interactions were set to zero, and no loop-extruders or plectonemes were included in the simulations. The models were then sampled for contact probabilities as before.

For the experimental data from Bintu *et al.*, we required estimates for the shape and size of the confinement volume, the monomer length, $b$, and the number of monomers each bin is mapped to, $n$. We chose to consider a spherical volume of confinement, with a radius $R$ given by one half of the mean maximal cross-section of the chromatin region in the chromatin tracing data. For each data set, we calculated the distribution for the separation $d$ between neighboring 30 kb regions, and used $\langle d^2 \rangle = nb^2$ to set $b$. We found that the choice of $n$ did not significantly affect our results (Supplemental Figure S9). Unless otherwise stated, results are shown for $n = 10$. We simulated confined random walks, and tracked how frequently every $n$th monomer was within a distance $< 150$ nm of each other. Code is available in [5].

#### S7 $P_3(s)$ curves

Given a three-point contact frequency array $M$, we calculated $P_3(s)$ curves as follows. For each three-point contact $i < j < k$, add $M(i, j, k)$ to $P_3(k - i)$. Divide each $P_3(s)$ by the number of three-point contacts $i < j < k$ with $k - i = s$.

We note that for a circular chromosome of length $N$, the genomic length of the largest loop is given by either $s = k - i$ or $s = N - k + i$. For consistency with 2D averaged plots, we chose $s$ equal to the sum of the two smaller loops in the three-point contacts, so that lines of $x + y = s$ on the 2D averaged plots still correspond to single points on the $P_3(s)$ curves. Using the minimum genomic length would map points with $N/2 < s < 2N/3$ to $N - s$. We hence display the plots only for $s < N/2$. As long as the same definition is used for comparing predicted $P_3(s)$ curves, the comparison is informative.

#### S8 $P_3(s)$ curves and averaging over sub-loop sizes

Just as a $P(s)$ curve contains less information than the Hi-C map from which it is derived, two models with different three-point contact statistics could still have similar $P_3(s)$ curves. To assess whether error cancellation when averaging over three-point contacts containing loops of different sizes results in improved $P_3(s)$ curves, we examine plots where the $x$ and $y$ axes correspond to the genomic lengths of the two shortest loops in a three-point contact.

Given a three-dimensional array $M$ corresponding to three-point contact data, we defined 2D averaged plots $A$ (Supplemental Figure S2) as follows. For a three-point contact $i < j < k$, let $x = \max(j - i, k - j)$, $y = \min(j - i, k - j)$. Increment $A(x, y)$ with $M(i, j, k)$. Divide each $A(x, y)$ by the number of three-point contacts with $x = \max(j - i, k - j)$ and $y = \min(j - i, k - j)$.

For a circular chromosome of length $N$, the algorithm must be adjusted. We calculated the minimum genomic distance between each pair of points in the three-point contact, $\min(j-i, N-j+i)$. $x$ and $y$ were chosen as the two lowest minimum genomic distances.

For all formulae the deviation of simulated data from predictions is largely a function of $s = x + y$. This means that $P_3(s)$ curves capture three-point contact probabilities averaged over genomic location.

### S9 Statistical analysis

Z-scores and two-tailed p-values for three-point contact frequencies were calculated by presuming that the number $k$ of three-point contacts observed in $n$ samples follows a binomial distribution $k \sim B(n, p)$, where $p$ is the predicted frequency of the three-point contact. Values were calculated using the HypothesisTests package for Julia [9].

The Benjamini-Hochberg procedure [10, 11] was used to analyze whether deviations were significant when the large number of possible three-point contacts was taken into account. The Benjamini-Hochberg procedure controls the false discovery rate (FDR), or the probability that we falsely reject the hypothesis for an individual three-point contact. If an FDR of $\alpha$ is required, all hypotheses with an adjusted p-value smaller than $\alpha$ should be neglected. Adjusted p-values (Supplemental Figures S4, S6) were calculated using the MultipleTesting package for Julia [12].

### S10 Algorithm by Liu *et al.* for Bintu *et al.* data

The code published by [13] was used to analyse the Bintu *et al.* data (Supplemental Figure S8). The script for Tri-C data was adapted to take in the Bintu *et al.* data as input, and to calculate a prediction for three-point contacts around every experimentally tracked locus. The results were combined into a 3D three-point contact probability array. The 3D array was analyzed in the same way as for the ideal polymer, loop-extruder, and pairwise interaction approximations, after scaling the results for each viewpoint to match the experimentally observed three-point contact counts.

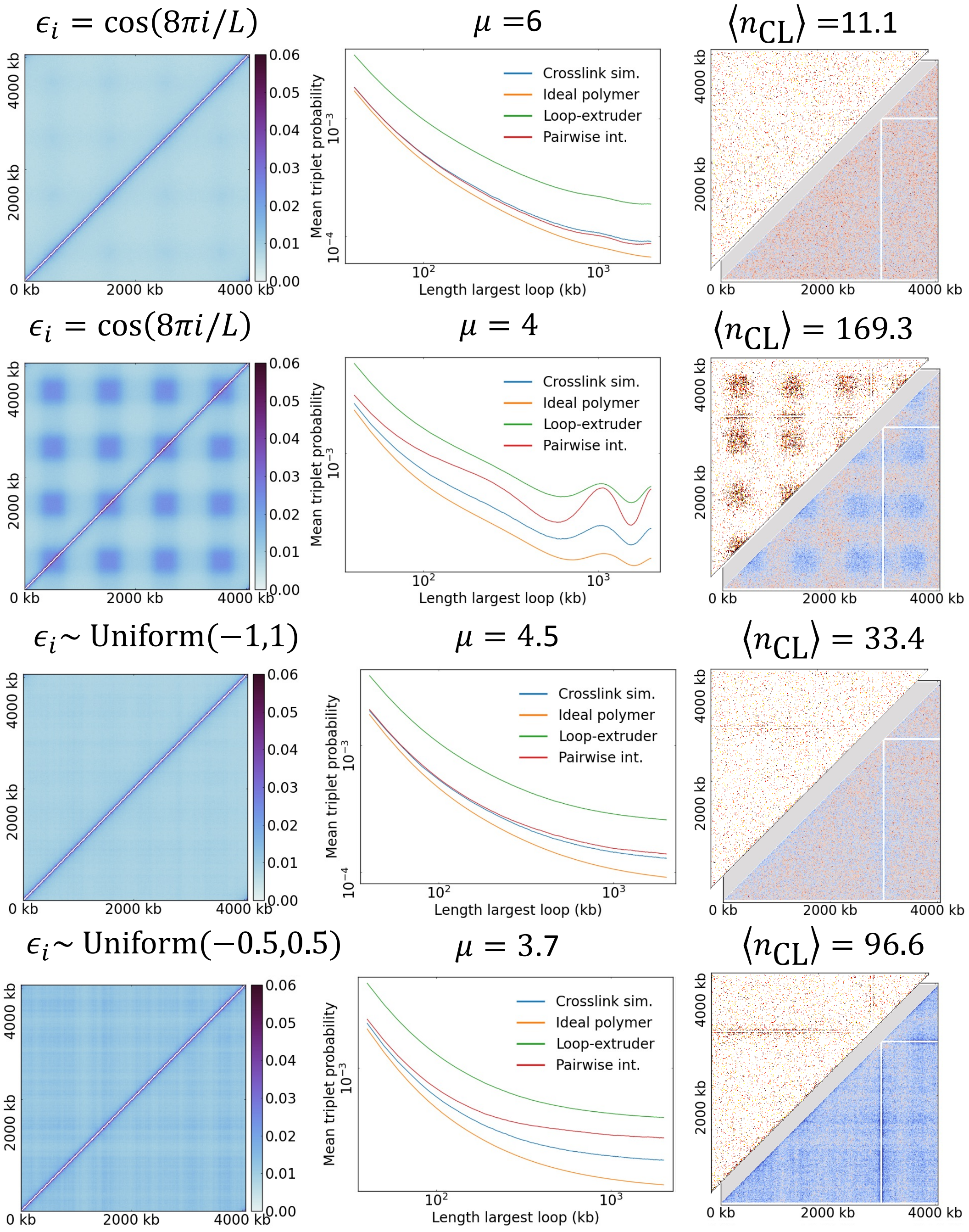


Figure S1: **Different crosslinker affinities** Each row corresponds to a different set of cross-linking affinities $\epsilon_i$ and chemical potentials $\mu$. The expected number of cross-linkers $\langle n_{\mathrm{CL}} \rangle$ for each potential is also included. Whether a periodic or random set of affinities is used, the pairwise interaction potential performs the best until the number of cross-linkers is of order 100. At this point the pairwise interaction formula starts to give a significant over-estimate, but it still performs better than the loop-extruder approximation.

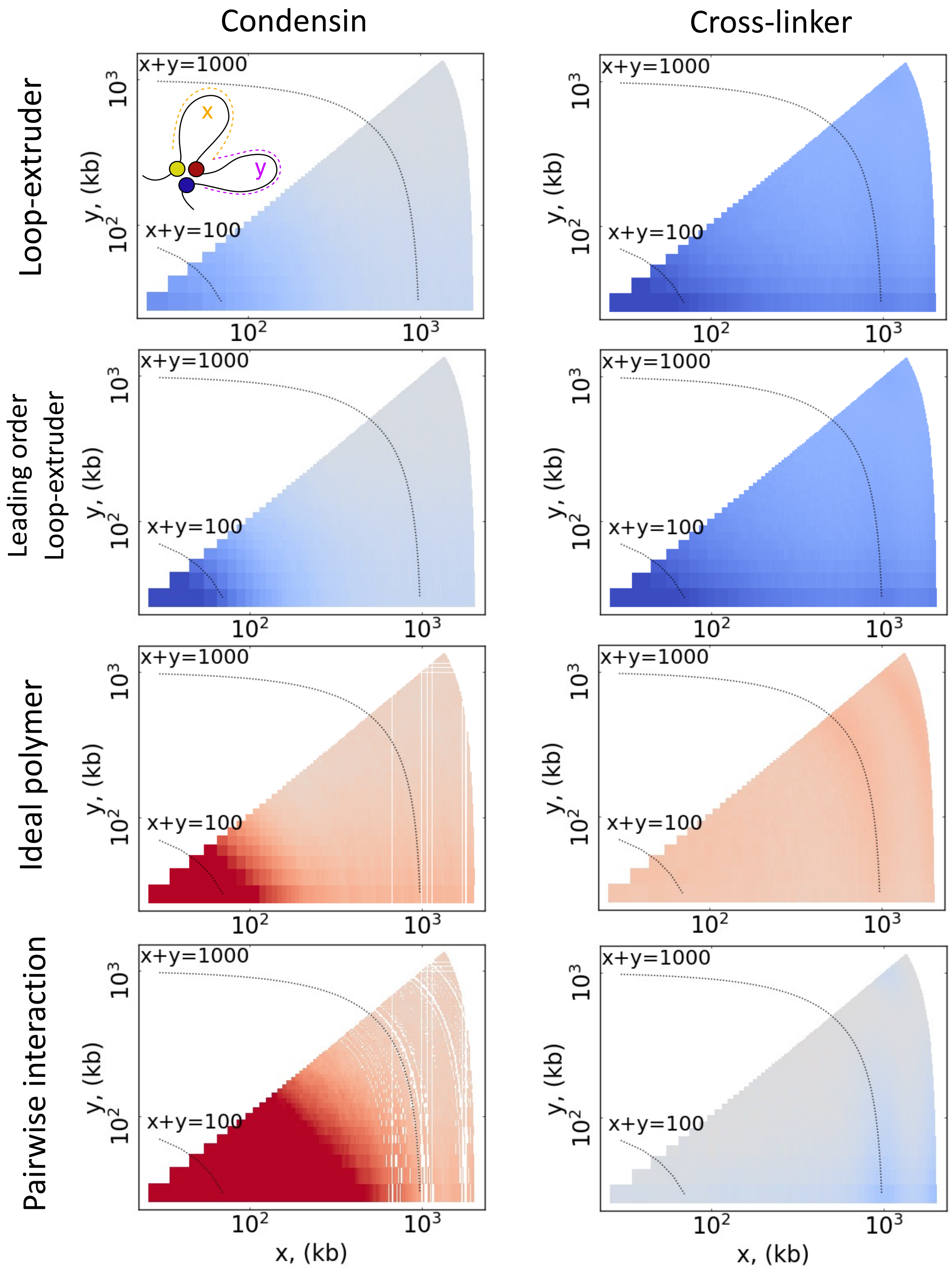


Figure S2: **2D averages of z-scores for simulated data** A table of averaged z-scores for different predictions for three-point contacts sampled from the cross-linker and condensin simulations. The intensity at $x > y$ is the average z-score for all three-point contacts where the two shorter loops are of lengths $x$ and $y$. Simulation parameters as in [Fig.2].

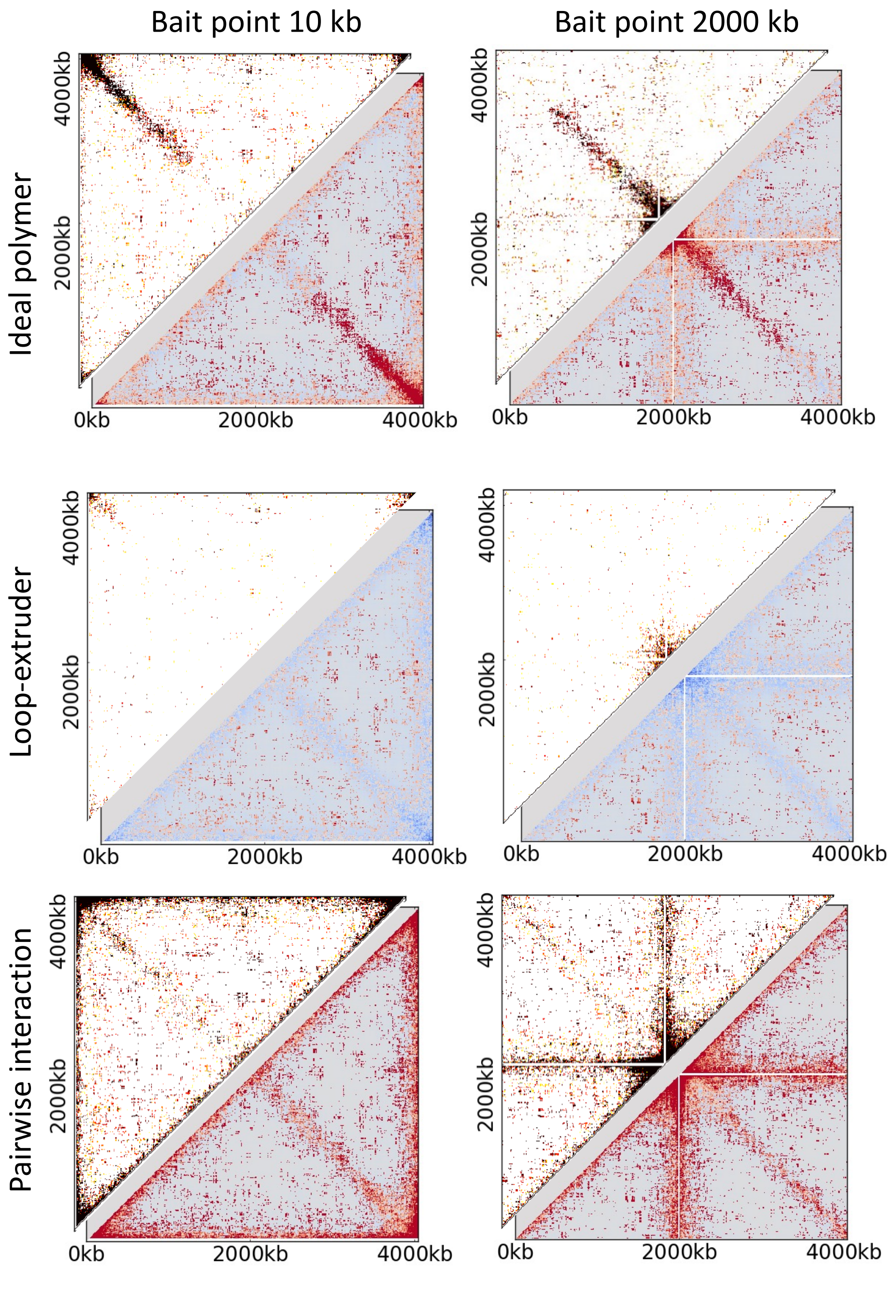


Figure S3: **Effect of bait point for simulation results.** Tables show p-value and z-score maps for two different bait points for the condensin simulations, similar to [Fig.3(c)] and using the same parameters.

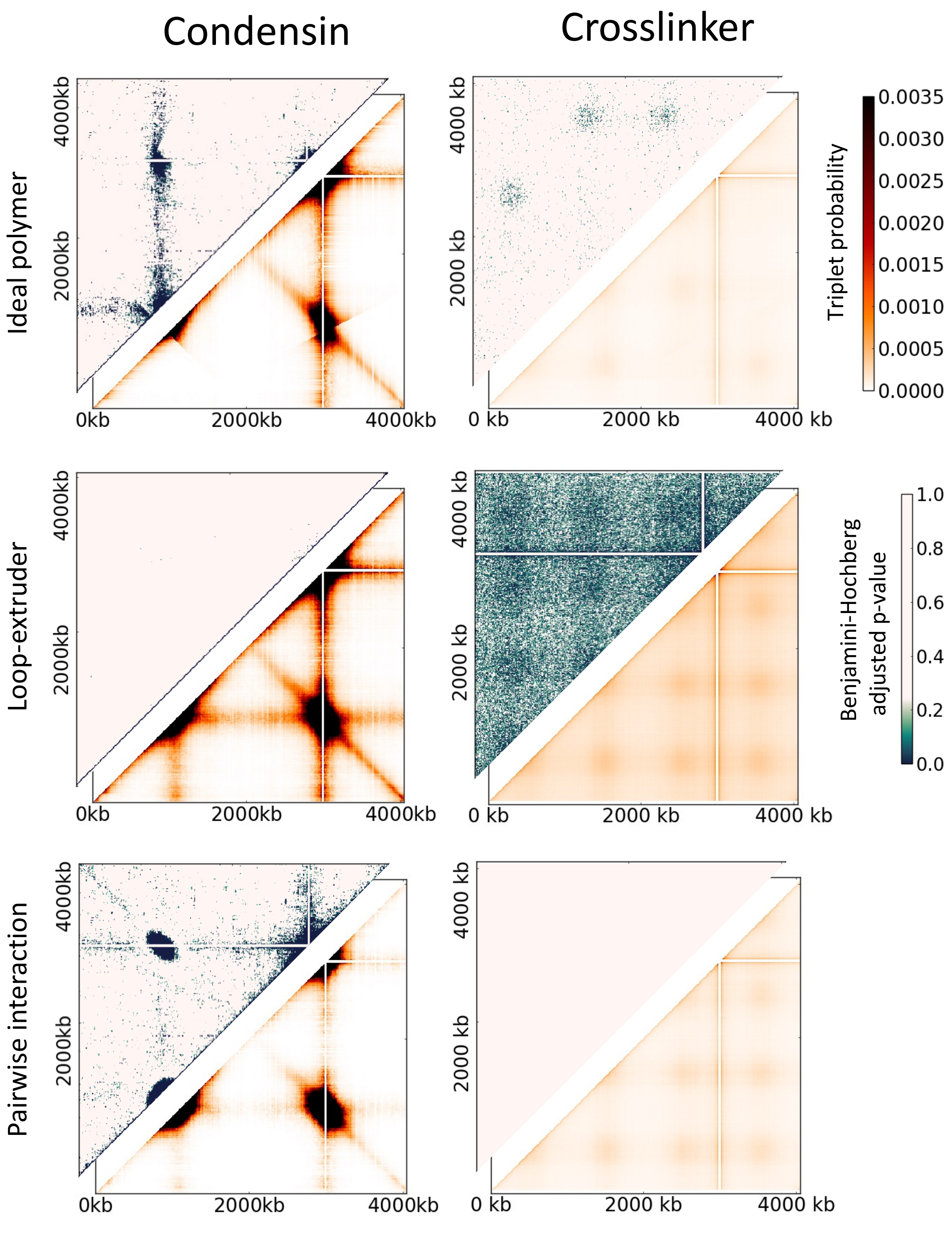


Figure S4: **Benjamini-Hochberg p-values for simulated data.** Table shows three-point contact probability maps (bottom-right) and Benjamini-Hochberg adjusted p-values for data from the crosslinker and condensin. Simulation parameters as in [Fig.2].

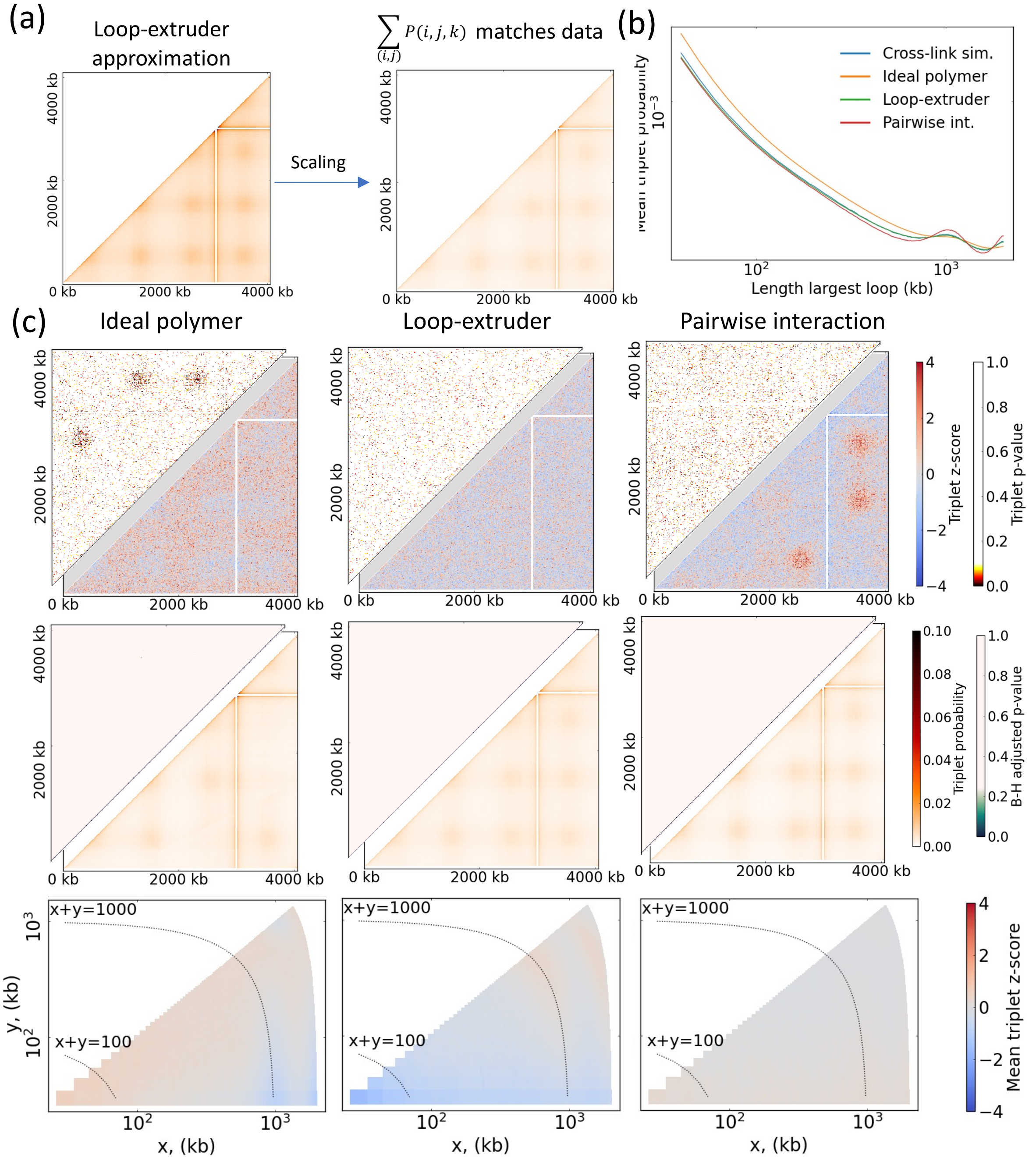


Figure S5: **Scaling of predictions for cross-linker simulations.** **(a)** Illustration of the effects of scaling the loop-extruder prediction for three-point contact data displayed in [Fig.3(a)]. Each predicted three-point contact probability is multiplied by the sum over the observed probabilities divided by the sum over predicted probabilities. **(b)** $P_3(s)$ curves similar to Figure 3A,B for scaled predictions. **(c)** P-value/z-score, Benjamini-Hochberg/three-point contact probability, and average z-score plots for scaled predictions of cross-linker three-point contact data.

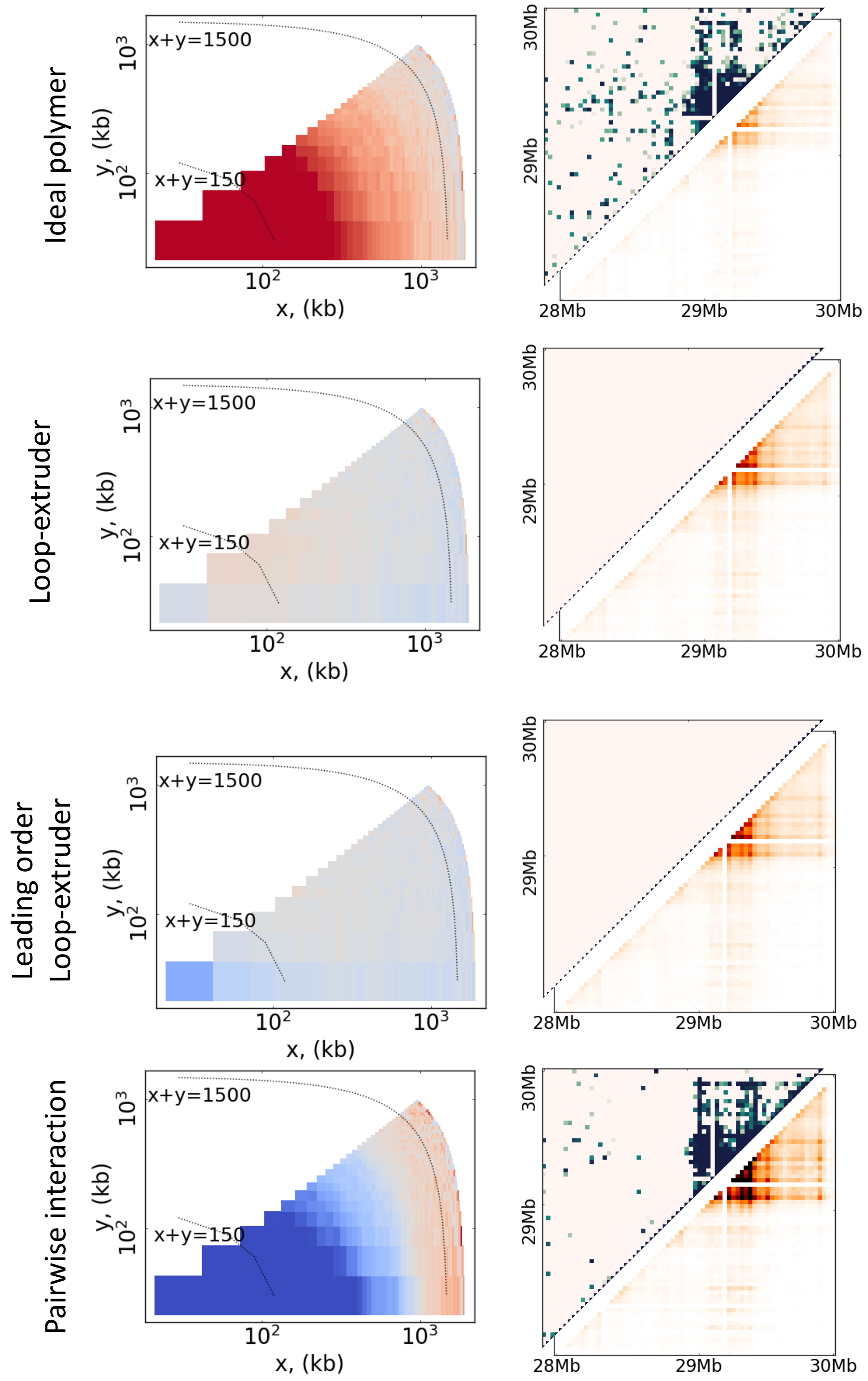


Figure S6: **Benjamini-Hochberg p-values and 2D averaged z-scores for Bintu *et al.* data.** Plots and color-scales are defined in Supplemental S2 and S4.

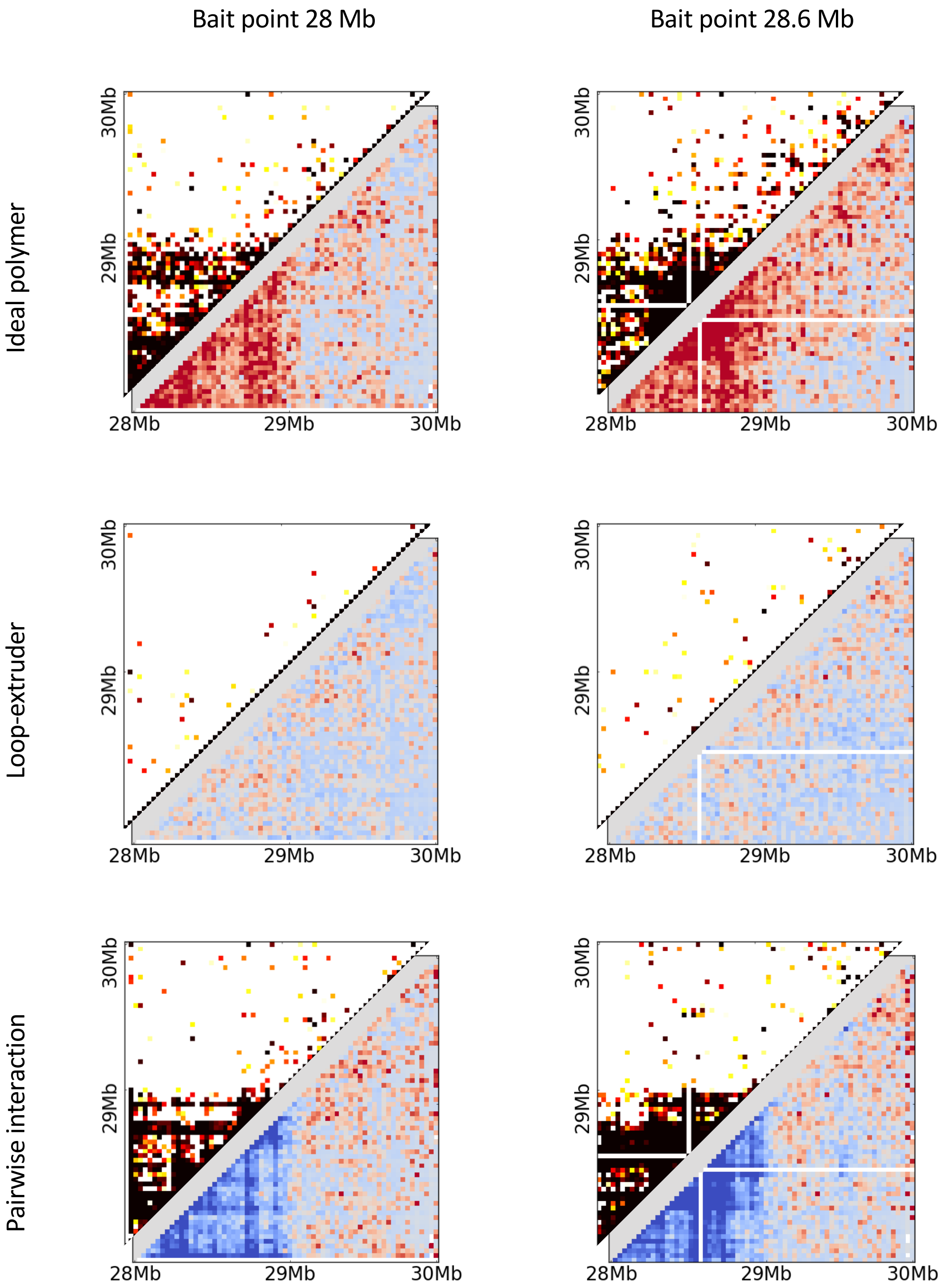


Figure S7: **Effect of bait point for Bintu *et al.* data.** Table shows similar plots to [Fig.4(d)] for two different bait points on the chromosome.

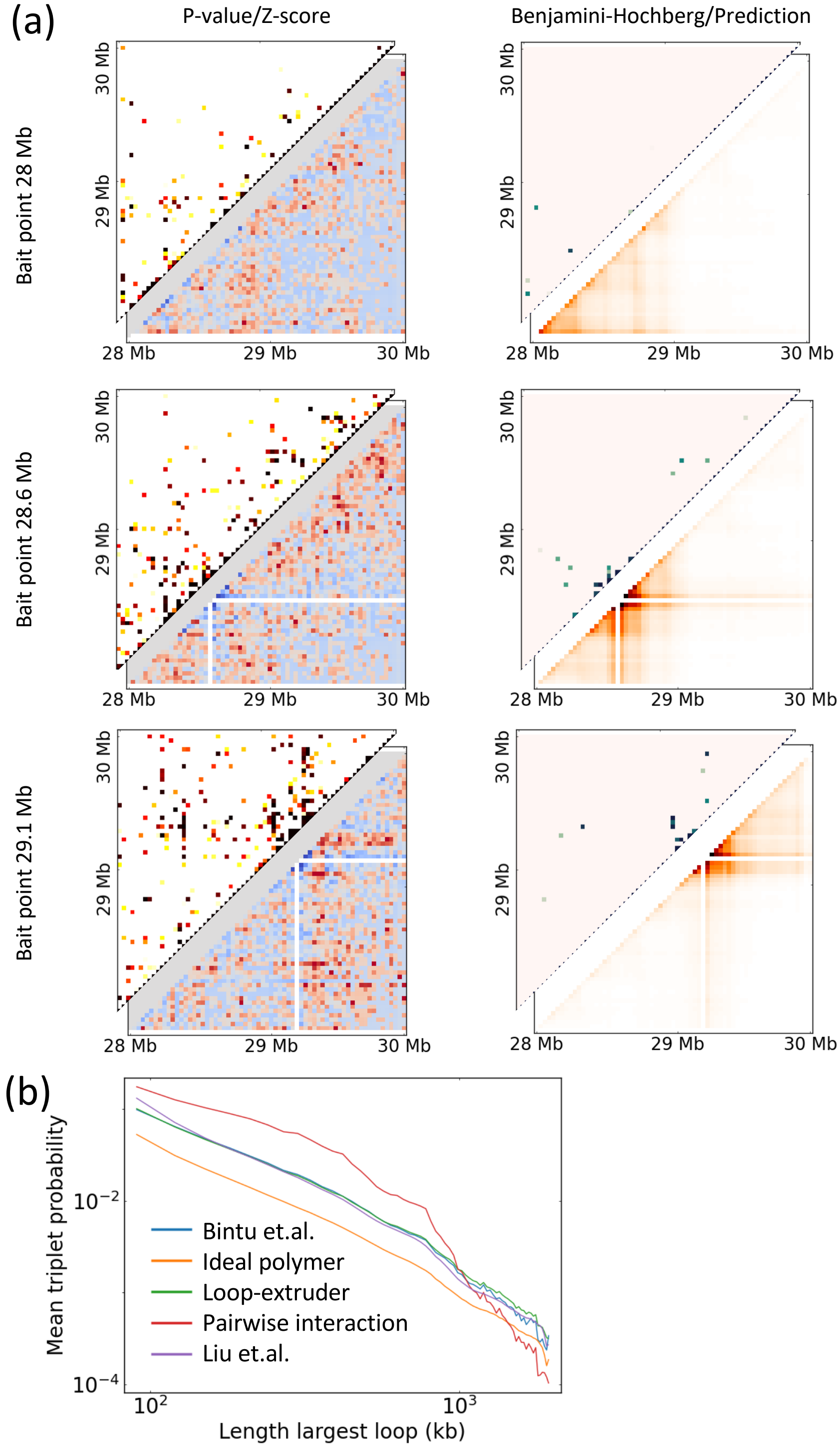


Figure S8: **Liu *et al.* prediction for Bintu *et al.* data. (a)** Comparison of Bintu *et al.* data to the prediction constructed following [13]. **(b)** $P_3(s)$ curve for prediction following [13].

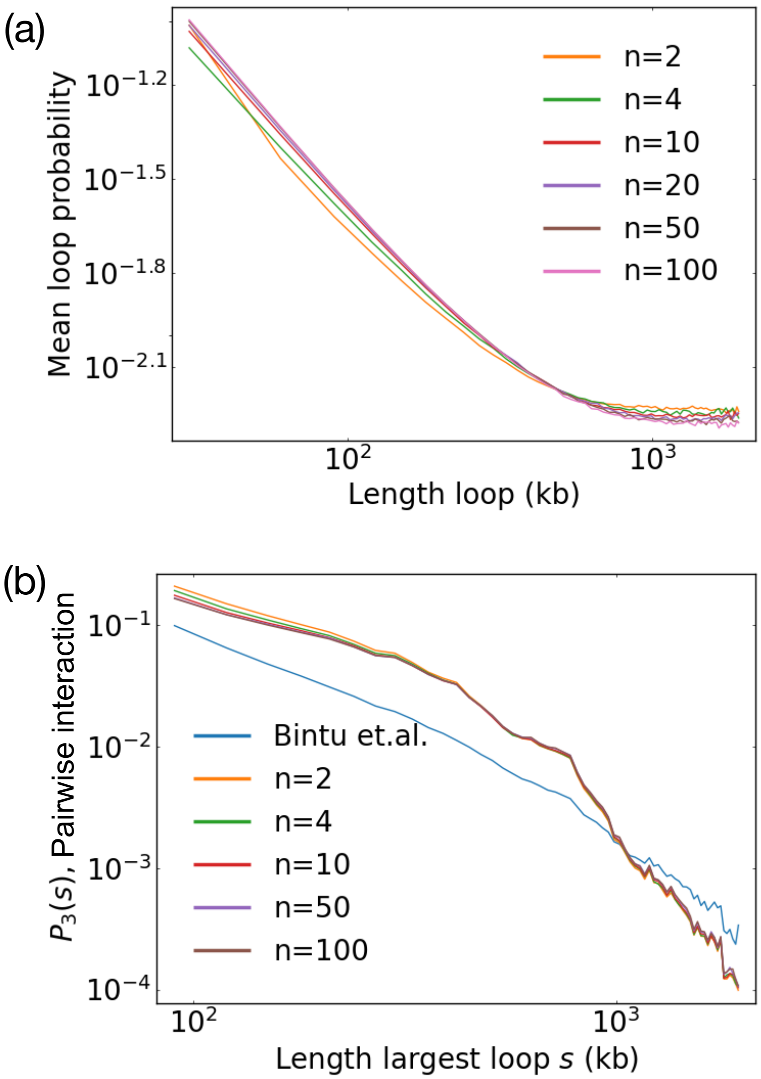


Figure S9: **Effect of $n$ on pairwise interaction formula is small.** **(a)** $P(s)$ curves for simulations of a non-interacting polymer in confinement, with different numbers of monomers corresponding to 30 kb segments. **(b)** $P_3(s)$ curve predictions of the pairwise interaction approximation, found by using the $P_0(n,m)$ distributions corresponding to different values of $n$.
